# Beyond a single patch: local and regional processes explain diversity patterns in a seagrass epifaunal metacommunity

**DOI:** 10.1101/482406

**Authors:** Keila A Stark, Patrick L Thompson, Jennifer Yakimishyn, Lynn Lee, Emily M Adamczyk, Margot Hessing-Lewis, Mary I O’Connor

## Abstract

Ecological communities are jointly structured by dispersal, density-independent responses to environmental conditions and density-dependent biotic interactions. Metacommunity ecology provides a framework for understanding how these processes combine to determine community composition among local sites that are regionally connected through dispersal. In 17 temperate seagrass meadows along the British Columbia coast, we tested the hypothesis that eelgrass (*Zostera marina* L.) epifaunal invertebrate assemblages are influenced by local environmental conditions, but that high dispersal rates at larger spatial scales dampen effects of environmental differences. We used hierarchical joint species distribution modelling to understand the contribution of environmental conditions, spatial distance between meadows, and species co-occurrences to epifaunal invertebrate abundance and distribution across the region. We found that patterns of taxonomic compositional similarity among meadows were inconsistent with dispersal limitation and meadows in the same region were often no more similar to each other than meadows over 1000 km away. Abiotic environmental conditions (temperature, dissolved oxygen) explained a small fraction of variation in taxonomic abundances patterns across the region. We found novel co-occurrence patterns among taxa that could not be explained by shared responses to environmental gradients, suggesting the possibility that interspecific interactions influence seagrass invertebrate abundance and distribution. Our results add to mounting evidence that suggests that the biodiversity and ecosystem functions provided by seagrass meadows reflect ecological processes occurring both within meadows and across seascapes, and suggest that management of eelgrass habitat for biodiversity may be most effective when both local and regional processes are considered.

## Introduction

Understanding how local environmental conditions, regional connectivity by dispersal and biotic interactions jointly structure the composition of communities is a central challenge in ecology (Ricklefs and Schluter 2003, Vellend 2010, Leibold and Chase 2017). Metacommunity ecology (Leibold et al. 2004, Leibold and Chase 2017, Thompson et al. 2020) offers a framework for understanding community assembly processes across spatial scales. In recent years, applications of the metacommunity framework have emphasized the underlying processes that give rise to abundance and diversity patterns (Brown et al. 2017, Leibold and Chase 2018, Thompson et al 2020). Thompson et al. (2020) framed the metacommunity concept based on three fundamental processes that together govern the dynamics of populations and communities: 1) Density-independent responses to environmental conditions, 2) Density-dependent biotic interactions (ie. inter and intra-specific competition, predation or facilitation influencing population growth and co-existence), and 3) Dispersal influencing connectivity across a landscape or seascape. We apply this framework to understand the contribution of these processes to a seagrass-associated invertebrate metacommunity.

Many coastal environments that host high biodiversity occupy spatially structured habitats (e.g. coral reefs, kelp forests, seagrass meadows), and many marine species have dispersing life histories that link populations in distinct habitat patches. The importance of interspecific interactions (Berlow 1999, Sala and Graham 2002), and dispersal-driven population dynamics (Levin and Paine 1974, Gaines and Roughgarden 1985) in coastal and marine communities has long been recognized. These combined roles of dispersal, the local environment, and biotic interactions suggest patterns of marine biodiversity reflect metacommunity processes (Boström et al. 2006).

Seagrasses are foundation species that support high productivity and faunal diversity (Orth et al. 1984, Duffy et al. 2015). They form meadows separated by deeper water, un-vegetated seafloor, or other vegetated habitats, and associated fauna disperse among meadows (Boström et al. 2006, 2011). Diverse invertebrate assemblages including snails, amphipods, isopods, and polychaete worms live among seagrass leaves, providing food sources for larger invertebrates, fish, and birds (Best and Stachowicz 2014, Huang et al. 2015). The grazers within this group consume detritus, macroalgae, or the seagrass itself (Valentine and Heck 1999), and many taxa exert top-down control on epiphytic microalgae that compete with the seagrass for light and nutrients (Sand-Jensen 1977, Duffy and Stachowicz 2006). Seagrass-associated epifauna exhibit a range of dispersal modes (fast swimming by isopods; slow crawling by snails; permanent attachment in bivalves) and reproductive strategies (brooding versus broadcast spawning) which influence dispersal rates and the distances over which meadows are demographically connected.

Seagrass meadows occur in a wide range of temperature, salinity and hydrodynamic environments, such that meadows may differ in their suitability for various invertebrate taxa (Yamada et al. 2007, 2014). Evidence for the importance of environmental filtering -- or the tendency for taxa to exist where local environmental conditions are ideal -- has been reported in several seagrass-associated fish and invertebrate systems (Baden et al. 2010, Robinson et al. 2011, Iacarella et al. 2018).

Though interspecific interactions among seagrass-associated invertebrates are not extensively documented, there is some evidence of competition and predation influencing community structure (Nelson 1979, Best and Stachowicz 2014). Predation can reduce abundances of vulnerable invertebrate taxa, allowing others to increase in abundance (Nelson 1979, Baden et al. 2010, Best and Stachowicz 2014, Amundrud et al. 2015, Huang et al. 2015). Despite some evidence for competition among epiphytic grazers for shared food sources (Edgar 1990, Bruno and O’Connor 2005) there is no evidence of competitive dominance to the point of exclusion in the field (Nelson 1979, Best and Stachowicz 2014).

While research in past decades has found evidence for direct and indirect influences of local characteristics such as habitat complexity (Orth et al. 1984), primary biomass (Cébrian and Duarte 1998, Gullström et al. 2012), and nutrient availability (Virnstein and Howard 1987) on faunal diversity and abundance, recent studies have focused more on regional scale or multi-meadow processes such as dispersal (Whippo et al. 2018, Lefcheck et al. 2016, Stier et al. 2019, Yeager et al. 2019), and have found that the influence of local-scale factors are often overriden by regional patterns of inter-meadow connectivity (Lefcheck et al. 2016, Stier et al. 2019, Yeager et al. 2019). This pattern has been attributed to relatively rapid life histories and high inter-meadow dispersal in seagrass-associated organisms (Lefcheck et al. 2016).

We designed this study to better understand the contributions of dispersal, local environmental conditions, and interspecific interactions with the largest spatially explicit (in geographic extent) dataset of seagrass metacommunity diversity. Drawing upon metacommunity theory, we tested the hypotheses that: (H1) Spatial distance *per se* within a region does not confer community dissimilarity because meadows are well-connected by dispersal, even at larger spatial scales (*dispersal limitation hypothesis*); (H2) Differences in local-scale environmental factors such habitat structure, temperature, and salinity drive some differences in invertebrate presence and abundance owing to differences in environmental niches among taxa, however the contribution of environmental factors is smaller than the contribution of spatial distance/ dispersal (*environmental filtering hypothesis*); (H3) No clear patterns of spatial co-occurrence among epifaunal species emerge because high dispersal and environmental responses override signatures of biotic interactions (predation, competition) that would produce such patterns (*biotic interactions hypothesis*).

## Methods

### Study sites

We sampled epifaunal invertebrate diversity in 17 *Zostera marina* (L.) meadows spanning the entire coast of British Columbia (approximately 1000 km distance) from late June through August of 2017. Within five regions that each contain numerous eelgrass meadows, we sampled a few meadows for our study (Fig. 1a): Haida Gwaii (two meadows); the central coast of BC near Calvert Island (four meadows); southern Clayoquot Sound (three meadows); Barkley Sound on the West coast of Vancouver Island (three meadows); the Southern Gulf Islands (five meadows). Meadows varied in environmental conditions, such as seagrass shoot size and density, hydrological regimes, and freshwater outflow influencing salinity.

**Fig. 1A.**
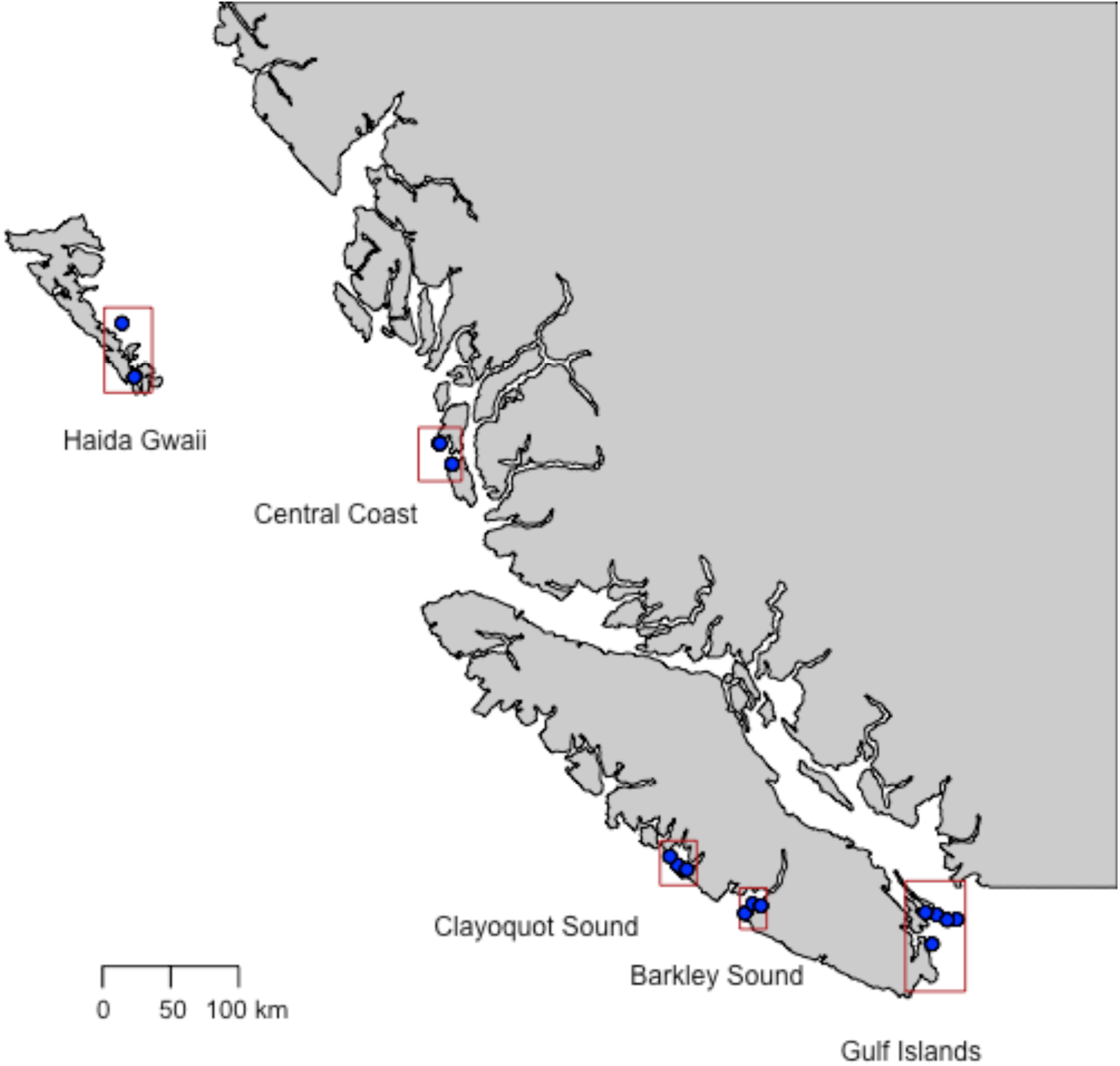
Map of coastal British Columbia showing eelgrass meadow sites in Haida Gwaii (HL, RA), Clayoquot Sound (DK, EB, IN), Barkley sound (SA, DC, RB), near Calvert island in the central coast (TB, TN, CS, CI), and the southern gulf islands (GB, JB, CB, LH, SS).

### Field sampling

We sampled three subtidal meadows with SCUBA in the Barkley Sound (SA, DC, RB) and Central Coast (TB, TB, CI, CS) regions. All others were sampled by wading or snorkeling at low tide. We conducted collections within a six-week period of peak seagrass growth. We used 0.25 m x 0.25 m quadrats to collect all above-ground eelgrass, epifaunal invertebrates, eelgrass detritus, and macroalgae in each sample, following Whalen et al. (2013). Six quadrats were arranged in a 15 m x 30 m array (Appendix 1: Fig. S1) in the middle of the meadow to avoid edge effects. We uprooted eelgrass shoots at the first node, leaving rhizomes to avoid sampling infauna that live in the sediments, removed all other above-ground biomass (detritus, macroalgae, associated epifauna) within each quadrat by hand, and immediately placed all the contents into a 300 µm mesh or plastic Ziploc bag for transport to the lab.

### Environmental data

We acquired water quality data from regional data sources. The abiotic water quality data were annual means pulled from Bio-ORACLE (Assis et al. 2018), including oxygen, nitrates, phosphates, silicates, salinity, maximum current velocity and sea surface temperature. Bio-ORACLE is a database of marine environmental layers gathered from several satellite and in-situ sources at a spatial resolution of 0.08°, and has been shown to accurately model distributions of shallow-water invertebrate species (Tyberghein et al. 2012, Assis et al. 2018). Sea surface temperature layers were taken from the Aqua-MODIS satellite, and values for nitrates, dissolved oxygen, and salinity were interpolated from in-situ measurements reported in the World Ocean Database (Tyberghein et al. 2012). All meadows were situated in distinct spatial cells, and thus had distinct values for the aforementioned variables. We excluded phosphate and silicate concentrations from our final analysis, as they strongly covaried with nitrates.

In each quadrat sample, we measured four biotic attributes summarizing habitat structure and food availability: eelgrass leaf area index (LAI), and eelgrass, eelgrass detritus, and algal dry mass. To quantify LAI, we measured leaf length, width, and blade number in five haphazardly-chosen shoots from each quadrat, and multiplied the average blade area per shoot by the quadrat-level shoot density. We dried eelgrass, detritus, and macroalgae in a desiccator oven (60° C for 48 hours) to measure ash-free dry mass.

### Invertebrate identification

Immediately after collection, we rinsed eelgrass shoots with fresh water, passing the water through a 500 μm sieve to remove epifaunal invertebrates. Invertebrates were preserved in 95% EtOH for identification with light microscopy (10x magnification). We identified invertebrates to the lowest taxonomic level possible using keys in the Light and Smith manual (Carlton 2007) and Kozloff (1996). In many cases, the lowest taxonomic resolution possible was family or genus, therefore our biodiversity survey likely underestimates full species diversity.

### Analysis

All statistical analyses were conducted in R (version 3.6.0; R Development Core Team 2019. We followed the Hierarchical Modelling of Species Communities (HMSC) framework (Ovaskainen et al. 2017) using the ‘Hmsc-R’ package (Tikhonov et al. 2019) to fit a hierarchical joint species distribution model (JDSM) with Bayesian inference. The framework uses traditional single-species distribution modelling by estimating species responses (presence or abundance) to environmental covariates across samples, but does so for all species simultaneously. It can use residual variation in occurrences to infer species co-occurrence patterns that do not result from shared responses to environmental covariates in the model (Ovaskainen et al. 2017). HMSC can also account for spatially hierarchical sampling structures. In these ways, the HMSC framework overcomes statistical limitations of previous methods used in metacommunity studies (Gilbert and Bennett 2010, Tuomisto et al. 2012, Brown et al. 2017).

Each individual species distribution model within the JDSM is a generalized linear model that describes the abundance of species *j* (where *j* = 1… *n*), where *y*_*ij*_ is the abundance of species *j* in sample *i, D* is the statistical distribution of the abundance data (Poisson distribution in this study), *L*_*ij*_ is the linear predictor to link species’ presence with environmental covariates, and *σ*^2^_*j*_ is a variance term for the abundance of species *j*: 

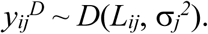

The linear predictor *L*_ij_ is described by fixed (*F*) and random (*R*) effect terms: 

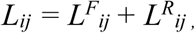

The fixed effect term *L*^*F*^_*ij*_ is modelled as a regression: 

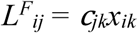

where *x*_*ik*_ represents the measured value for environmental covariate *k* for a given sample *i* (e.g., biomass in quadrat *i*), and parameter *β* represents the relationship between environmental covariate *k*, and the abundance of species *j.*

The random effect term *L*^*R*^_*ij*_, captures the variation in species abundances that cannot be explained by the measured covariates. It is further denoted as: 

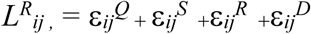

where the terms describe three random effects associated with our spatially nested sampling scheme (ε_*ij*_^*Q*^, ε_*ij*_^*S*^, ε_*ij*_^*R*^ for Quadrat (sample), Site (meadow), and Region), as well as a fourth spatially explicit random effect (spatial autocorrelation in species abundances). The spatially explicit random effect ε_*ij*_^*D*^ was calculated with latent factor analysis that takes into account spatial distance between all pairwise combinations of samples (see Ovaskainen et al. 2016a for details).

The three non-spatial random effect terms are assumed to have multivariate normal distributions ε_*ij*_^*Q*^ ∼ *N*(0, Ω^*Q*^), ε_*ij*_^*S*^ ∼ *N*(0, Ω^*S*^), ε_*ij*_^*R*^ ∼ *N*(0, Ω^*R*^). Variance terms Ω^*Q*^, Ω^*S*^, Ω^*R*^, are variance-covariance matrices (square matrices containing all taxa in the model), where the diagonal elements give species-specific residual variance in occurrences among samples, and the off-diagonal elements give residual co-variances between species pairs. The term “residual” refers to the fact that it is the variance unexplained by environmental covariates in the fixed effects predictor described above. These variance-covariance matrices (Ω^*Q*^, Ω^*S*^, Ω^*R*^, and Ω^*D*^) are parameters estimated using a latent variable approach described in Ovaskainen et al. (2016b). They were used to represent co-occurrence matrix R, where *j*_*1*_ and *j*_*2*_ refer to two species within the model, and 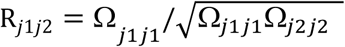. R describes the extent to which a given species pair co-occurs more positively or negatively than by chance, after controlling for possible shared responses to the same environmental covariates.

We estimated parameters with Markov chain Monte Carlo. Markov chains were run to 200 000 iterations, a burn-in length of 1000 iterations, and was thinned to retain every 10^th^ sample of the posterior distributions. We confirmed that Markov chains were well-mixed by visually inspecting trace plots. Estimates of parameter *β* (environmental responses across taxa) and Ω (variance-covariance matrices used to visualize species co-occurrence patterns independent of the environment) were extracted as 95% credible intervals. We evaluated the model fit (R^2^ and Root-Mean-Square Error or RMSE) using 4-fold cross-validation.

To test our three hypotheses, we visualized different aspects of the parametrized JDSM. To test for dispersal limitation (H1), we looked for increased pairwise community dissimilarity among samples with increased pairwise spatial distance. We predicted species abundances in every sample using our trained model, then calculated pairwise Bray-Curtis dissimilarity index (Bray and Curtis 1957) on these predicted abundances. We then conducted log-linear regression of predicted pairwise dissimilarity against pairwise spatial distance (km) on a logarithmic scale. Increased community dissimilarity could reflect dispersal limitation due to prohibitively far dispersal distances, *or* due to increasing differences in environmental conditions with increasing spatial scale. To distinguish between these possibilities, we compared a regression of predicted dissimilarity based on the full JDSM (including environmental covariates) with the predicted dissimilarity calculated from the same trained model, but with no effect of the environment (all environmental covariates were set to their mean values).

To test the environmental filtering hypothesis (H2), we analyzed *β* for each combination of species and environmental covariate to determine whether they had a positive or negative relationship. We also partitioned the variance in abundance explained by environmental covariates, spatial autocorrelation, and sampling design random effects.

To test the biotic interactions hypothesis (H3), we represented co-occurrence matrix R as a correlation plot for each random effect level, demonstrating the extent to which species pairs co-occurred more negatively or positively than predicted by their modelled correlation with environmental covariates at each spatial scale (Ovaskainen et al. 2017, Aivelo and Norberg 2018).

## Results

### Taxonomic abundance and distribution patterns

We identified 52 282 individuals representing at least 50 taxa across the region (Appendix 2: Table 2). Of these, 3% of individuals were bivalves, 12% were gastropods (snails and limpets), 14% were copepods, 41% were polychaetes (most of which were calcifying polychaetes *Spirorbis* sp.), 13% were gammarid amphipods, 6% were caprellid amphipods, and the remaining 11% included other crustaceans (isopods, tanaids, cumaceans, crabs, shrimp). These taxa span several phyla, diet types (herbivores, detritivores, suspension feeders), and dispersal strategies (brooding, broadcast spawning). Mean meadow-level taxonomic richness was 20 taxa per 0.0625 m^2^ of seagrass area.

Several taxa were present at all 17 meadows: harpacticoid copepods *Porcellidium* sp. (4% of all individuals across the entire region), snails *Lacuna* spp. (7%), and tanaids *Leptochelia* sp. (10%). We found genera *Mytilus* sp. (mussel) and *Nereis* sp. (polychaete worm) in all meadows, however we were not confident the same species were present at each meadow. Ten taxa were present in every region, but not necessarily every meadow: isopod *Pentidotea resecata*, gammarid amphipods *Monocorophium insidiosum, Ampithoe valida, Pontogeneia rostrata, Ampithoe dalli, Aoroides* spp., snails *Amphissa columbiana* and *Alia carinata*, and limpet *Lottia pelta*.

We excluded rare species (those found in fewer than 5% of samples, or fewer than 5 out of 102 quadrats) from our JDSM, leaving 33 taxa in our modelling analysis (Appendix 1: Table S2). This was necessary to avoid statistically over-inflating the importance of model covariates.

### Spatial connectivity via dispersal driving community similarity

The highest Bray-Curtis dissimilarity value was 0.99 for a pair of sites spaced 787 km apart (HL in Haida Gwaii and SA in Clayoquot Sound). The two communities at the highest pairwise distance (997 km, RA in Haida Gwaii and SS in the Gulf Islands) had a dissimilarity index of 0.8. The lowest dissimilarity index value was 0.59 at a pairwise spatial distance of 0.6 km (LH and SS in the Gulf Islands). A log-linear regression of Bray-Curtis dissimilarity calculated from predicted abundances using the JDSM showed that community dissimilarity increased with increased spatial distance (Fig. 1b, *y* = 0.75 + 0.023log(*x*), n = 272, R^2^ = 0.16). The maximum and minimim pairwise dissimilarity values were reduced when the effect of the environment was removed (*y* = 0.74 + 0.021log(*x*), n = 272, R^2^ = 0.07), suggesting environmental variables drive some dissimilarity in species composition with increased spatial distance.

**Figure 1b.**
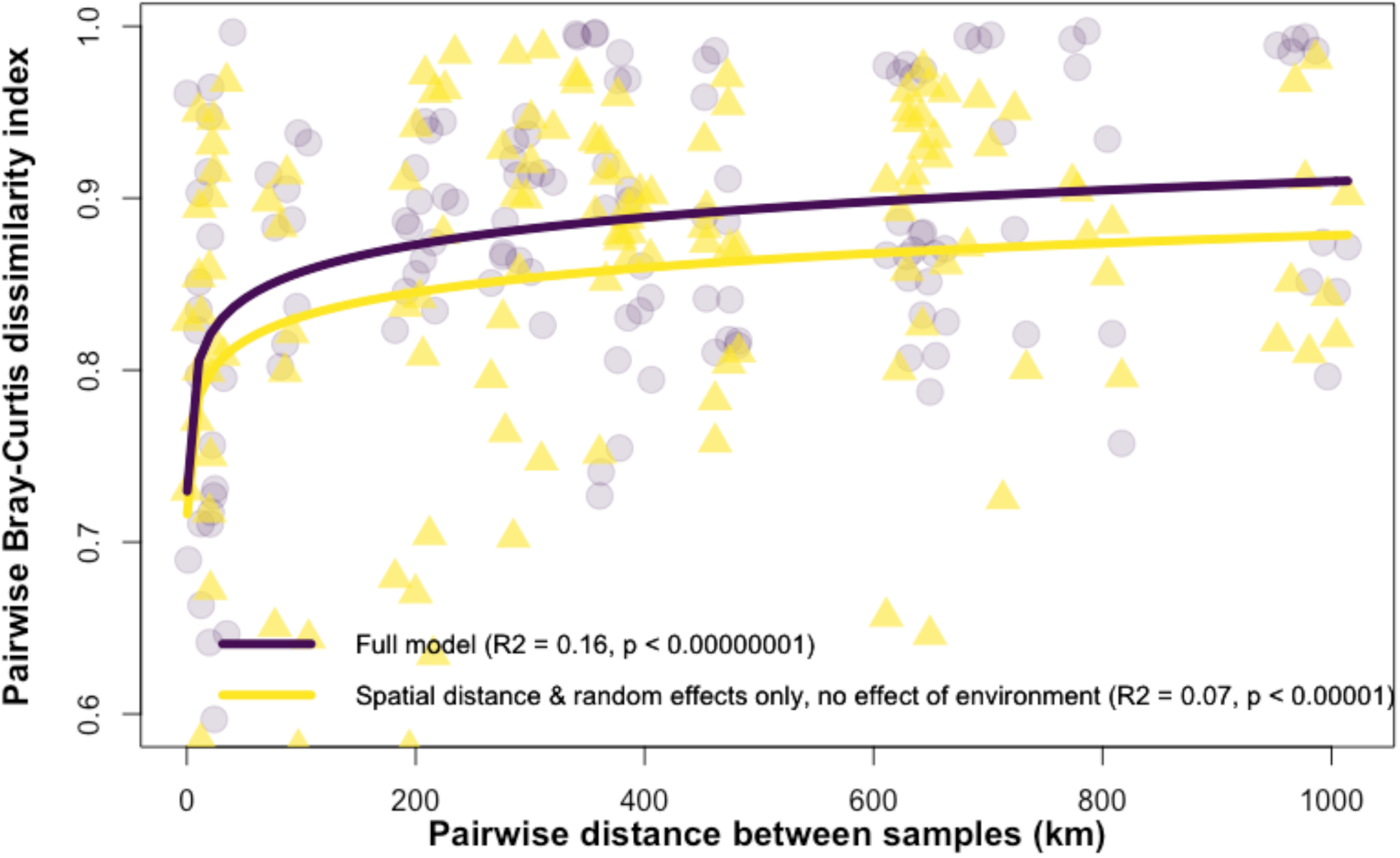
Bray-Curtis pairwise community dissimilarity index as a function of spatial distance (Euclidean) between every pairwise combination of meadows in our study. The purple points and line show pairwise dissimilarity predicted by the full joint species distribution model using the original environmental and spatial data. The yellow points and line show pairwise dissimilarity predicted by the same trained model, but the effect of all environmental variables has been removed (set to their mean values). The yellow points and line show calculated pairwise dissimilarity based on the raw abundance data.

### Environmental conditions

The assessment of model fit and variance partitioning analysis both showed that the importance of environmental covariates differed among taxa, and that overall the environment explained relatively a low proportion of variation in species abundances across the region (Fig 2a). While the mean R^2^ across individual species distribution models was 0.27, individual R^2^ values varied (0 to 0.68, Appendix 1: Table A2) indicating variability in model fit.

**Figure 2a.**
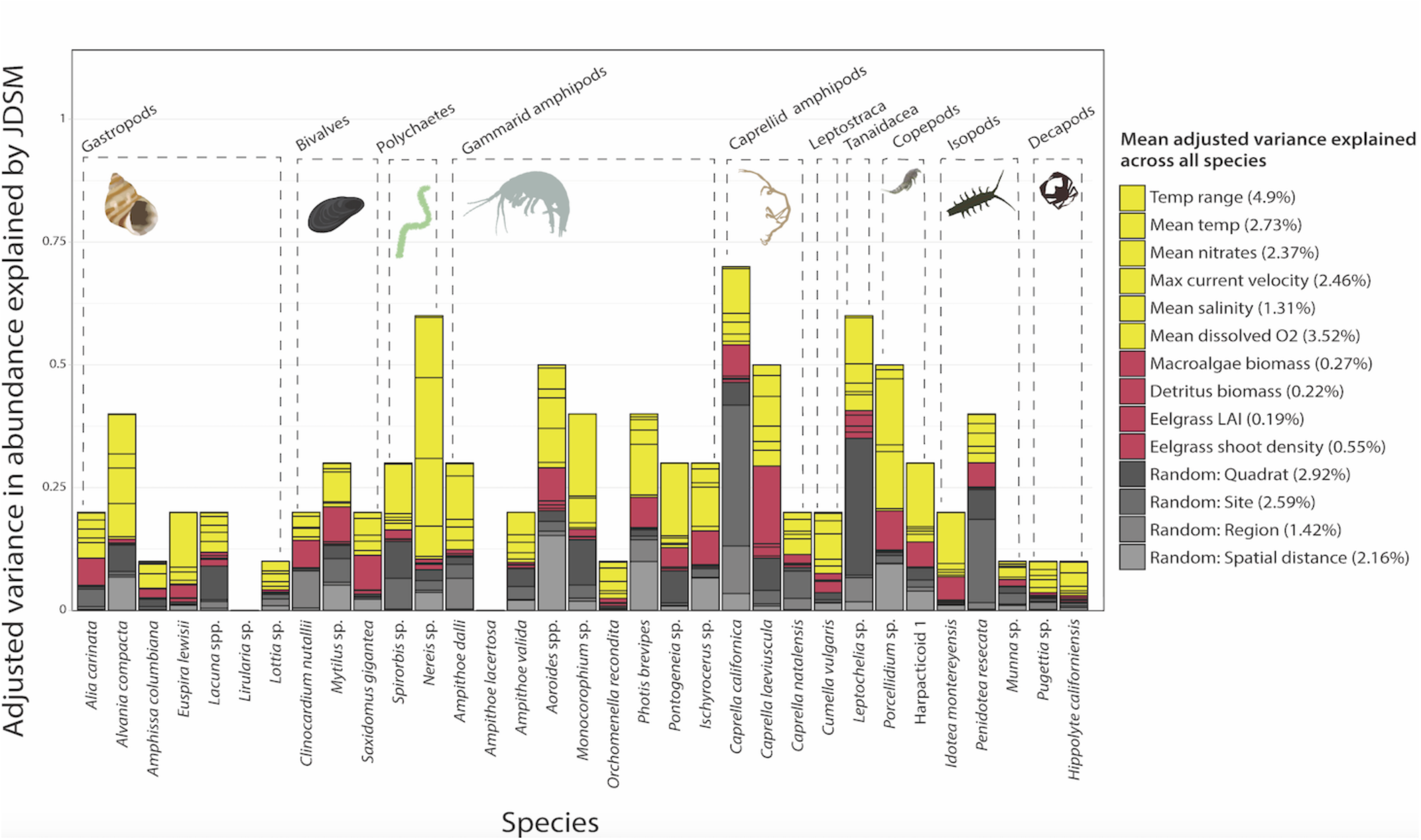
Variation partitioning of fixed and random effects within the joint species distribution model. Variances explained have been adjusted to reflect the model fit (Multiplied by Pseudo R2). An unadjusted version of the variance partitioning analysis can be found in Appendix 1, Fig. S3 of the Supplementary Information. Dashed lines indicate broad invertebrate taxonomic groupings. Grey cells represent the contribution of random effects associated with the sampling design. Yellow cells represent abiotic water quality covariates. Pink cells represent biotic covariates (food availability, habitat structure). The percentages next to the legend labels indicate mean variation explained by that covariate/ random effect across all species distribution models.

Overall, abiotic water quality variables explained more variation than biotic variables (Fig. 2a). Mean and range of sea surface temperatures and dissolved oxygen explaining the largest proportions of variation on average. Herbivore food availability (seagrass, algae biomass) and habitat structure (seagrass shoot density and LAI) had lower influence on abundance (pink bars, Fig. 2a). The importance of some environmental covariates differed markedly among taxa; for example, nitrate levels explained approximately 27% of variance in the abundance of gastropod *Alvania compacta* across meadows, but only 5% of variance in gastropod *Alia carinata* (Fig 2a). The mean variance in taxon abundance explained by spatial autocorrelation was 2.2% (pale gray bars, Fig. 2a), and other hierarchical random effects associated with the sampling design explained 2.92% (quadrat-level), 2.59 (meadow or site-level), and 1.42% (region-level) of variation.

The strength and directionality of responses to environmental variables also differed among taxa, based on inspection of the *β* parameters that estimate the relationships between environmental covariates and species abundances (Fig. 2b). None of the taxa in our study showed statistically supported responses (*β* within 95% credible interval) to all environmental variables. The few that showed a significant response to eelgrass LAI responded positively (higher LAI meant higher abundances). Thirteen taxa responded negatively to higher temperature ranges, and two responded positively (Fig. 2b). The majority of taxa (22 out of 33) had significant responses to dissolved oxygen levels. Of these, 8 responded negatively (high dissolved O_2_ yields lower abundances), and 14 responded positively (high dissolved O_2_ yields lower abundances).

**Figure 2b.**
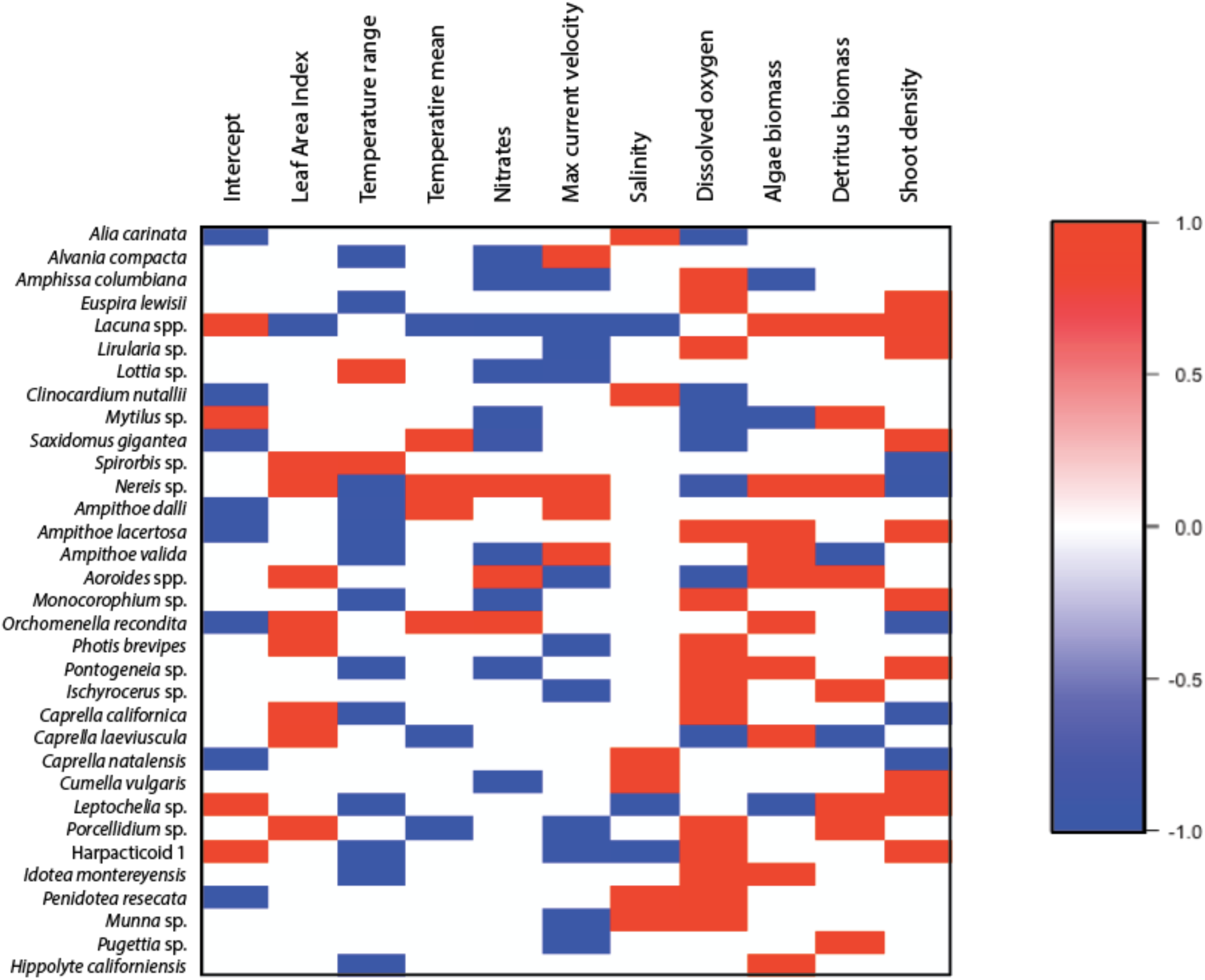
Heat plot summarizing beta (*β*) parameters, or responses to all environmental covariates in the joint species distribution model. Coloured cells indicate that the response is statistically supported (*β* falls within 95% credible interval). Red cells indicate positive responses to a given environmental covariate (higher abundance correspond with higher values of the environmental covariate), whereas blue cells indicate negative responses (lower abundance with higher values of the environmental covariate).

### Co-occurrence patterns

Co-occurrence patterns were less strong at the between-quadrat and between-region spatial scales (Fig 3a, 3c). In comparisons among quadrats, we observed more positive species co-occurrences than negative co-occurences (Fig. 3a). In comparisons among meadows, we identified, *post hoc*, two main species co-occurrence groupings (Fig. 3b). Members of the first group (hereafter referred to as *Leptochelia* group) included *Leptochelia* sp. (tanaid), *Photis brevipes* (gammaridean amphipod), *Amphissa columbiana* (snail), *Spirorbis* sp. (calcifying polychaete), *Nereis* sp. (polychaete worm), *Caprella laeviuscula* and *Caprella natalensis* (caprellid amphipods) (Fig. 3b). These taxa positively co-occurred more often than expected by chance or by environmental filtering. They negatively co-occurred with members of the second group, which included a harpacticoid copepod species, *Mytilus* sp. (mussel), *Porcellidium* sp. (copepod), *Ischyrocerus* sp. (gammarid amphipod), and *Penidotea resecata* (isopod) (hereafter referred to as harpacticoid group). *Alvania compacta* and *Lirularia* sp. (snails) positively co-occurred with each other but did not strongly co-occur with members of the two main groups. Remaining taxon pairs did not exhibit strong positive or negative co-occurrence patterns.

**Figure 3.**
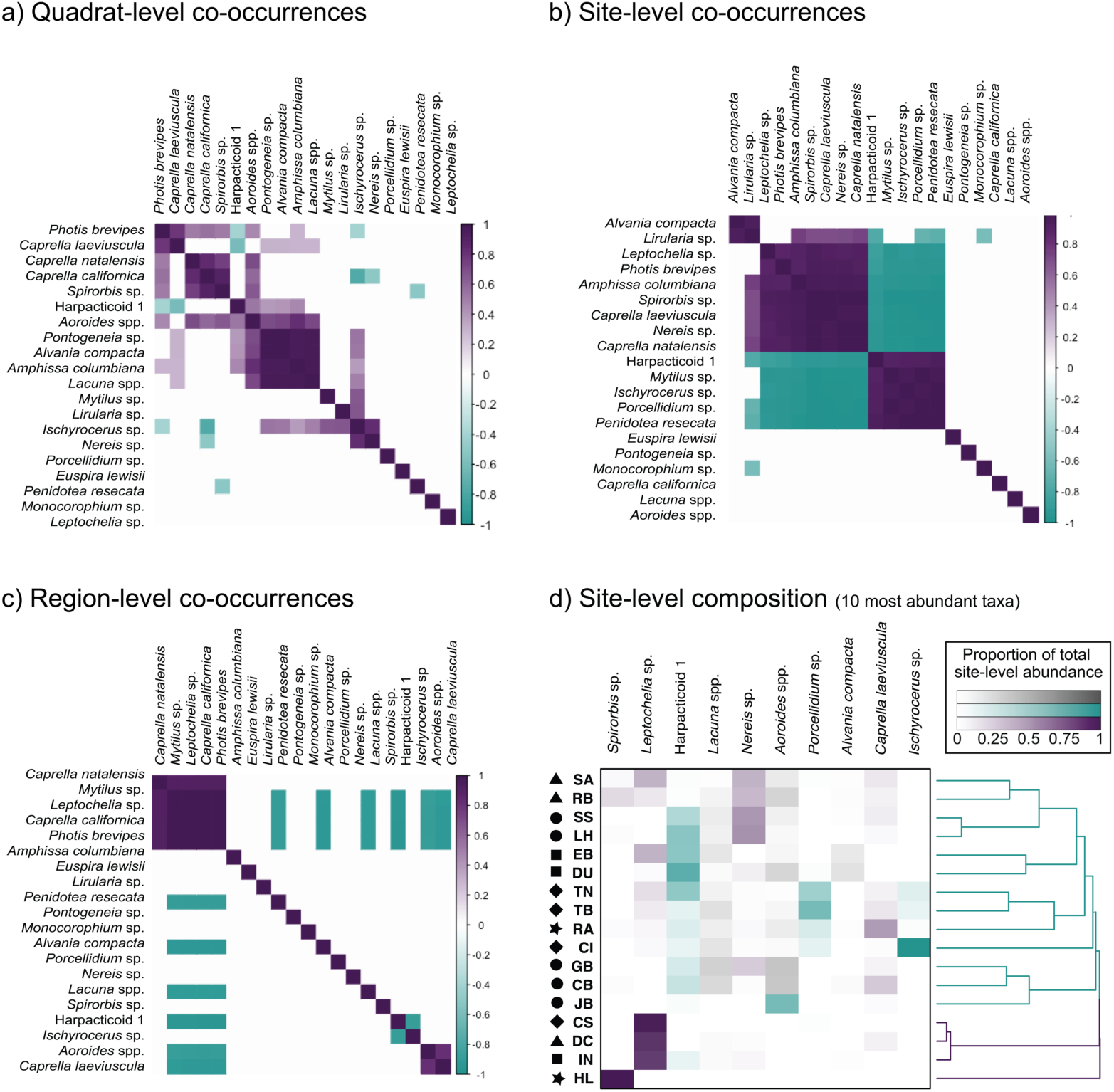
**a-c)** Correlation plots showing modelled a) quadrat-level, b) site-level, and c) region-level pairwise co-occurrences after removing the effect of shared responses to the environmental covariates in our model. Only the 20 most abundant species (according to total abundance across the metacommunity or study region) are represented. Purple cells represent positively co-occurring species pairs, and turquoise cells represent negatively co-occurring species pairs. Species names are ordered according to the output of hierarchical clustering with Ward’s criterion on pairwise co-occurrence values. **d)** Heatmap and cluster dendrogram depicting species relative abundance and compositional similarity across sites. Species are ordered by decreasing abundance from left to right. Cell colours correspond to the two main site-level co-occurrence groupings shown in Fig. 3b: purple or “*Leptochelia”* group, and blue or “Harpacticoid” group. Cell shade strength indicates proportional abundance at a given site (darker means higher relative abundance). The black symbols to the left of the site abbreviations indicate region membership; stars indicate Haida Gwaii sites, diamonds indicate Central Coast sites, triangles indicate Barkley Sound sites, squares indicate Clayoquot Sound sites, and triangles indicate Southern Gulf Islands sites.

The two co-occurrence groups were represented at all meadows, suggesting that members from these groups can be simultaneously present (Fig. 3d). However, their abundances tended to negatively co-vary; at meadows JB, HL, CB, RA, EB, DC, and DU, members of the *Leptochelia* group were most abundant, whereas Harpacticoid group members were most abundant at SS, SA, LH, GB and RB. Meadow HL was an outlier, as it was strongly dominated by *Spirorbis* sp. The emergent species groupings identified in Fig. 3b do not clearly correspond with geographical structure because the cluster analysis did not group meadows by region membership (black symbols, Fig. 3d).

## Discussion

With a biodiversity survey of 17 seagrass meadows across an approximately 1000km spatial extent, we fitted a joint species distribution model to test hypotheses about the contributions of dispersal, environmental filtering, and species interactions to epifaunal invertebrate abundance and distribution. We also documented at least two previously unrecognized species’ associations in the region. This study is the largest (in geographical extent) spatially explicit seagrass-based metacommunity study to our knowledge. We found little support for complete dispersal limitation even across over 1000 km of coastline, some support for the importance of environmental niche filtering, and co-occurrence patterns that might have arisen from interspecific interactions. Overall, these results suggest that seagrass-associated biodiversity patterns reflect ecological processes spanning local (meadow-scale) to regional scales, and provide additional support for recent findings that eelgrass-associated diversity reflects regional-scale ecological processes in addition to local scale processes (Lefcheck et al. 2016, Whippo et al. 2018, Stier et al. 2019, Yeager et al. 2019).

### Dispersal limitation is unlikely at this regional spatial scale

Our findings that: 1) Many species were present in all regions or all meadows; 2) The effect of spatial distance alone did not explain decay in community similarity (Fig. 1b); 3) There was a low contribution of spatial autocorrelation to variance in species abundance (Fig. 2a); and 4) Hierarchical cluster analysis did not group meadows according to spatial regions (black symbols in Fig. 3d), all suggest that distance between meadows likely does not confer dispersal limitation preventing populations at distant sites from exchanging individuals. It is however likely that meadows that are physically near each other likely experience higher rates of exchange in individuals than distant ones, given our observation that dissimilarity increases logarithmically with increased spatial distance. The six taxa that were present at all meadows and an additional ten taxa that were present in all regions had representation from across phyla and life history traits (including rare taxa), suggesting that no single taxonomic group or dispersal strategy had a consistently larger spatial distribution than others. Many meadows were small (< 1 ha), and epifaunal abundances can vary substantially within meadows from year to year (Nelson 1997, Douglass et al. 2010) such that local extinctions followed by rescue from nearby populations are likely (Thom et al. 1995). However, the specific pathway that dispersing organisms travelled to arrive at these meadows remains unknown. We used Euclidean distances in our analysis, which preserved the rank order of distances between meadows, but actual distances are likely greater due to oceanographic circulation patterns (Kinlan and Gaines 2003, Mitarai et al. 2008).

It is unclear whether distant meadows share taxa because they: 1) Are linked by direct dispersal via oceanographic processes (currents); 2) Are indirectly linked by dispersal via unsampled “stepping-stone” meadows or other habitat types; or 3) Were colonized by populations of the same species in a historical dispersal event, but have not seen the exchange of individuals since (Lefcheck et al. 2016). Seagrass-associated epifauna generally disperse passively between meadows, either through larval transport in currents or by “rafting” on floating pieces of seagrass or macroalgae. Rafts have been observed dispersing benthic and epifaunal invertebrates across phyla (echinoderms, peracarids, molluscs, annelids), and facilitate connectivity between coastal ecosystems across 100s of kilometres (Wichmann et al. 2012).

Our inferences are based on an analysis that necessarily emphasized common taxa – either numerically dominant or present in most meadows, or both. We observed seventeen rare taxa that were found in fewer than 5% of our samples, often at extremely low abundances (1-2 individuals per meadow, or per 6 quadrats) in our survey of over 50 000 individuals. The spatial extent of their range could be dispersal-limited for reasons other than prohibitively long travel distances. Populations with low abundances may not disperse in appreciable numbers, and therefore cannot establish populations as readily as abundant species; this is supported by our observation of a positive relationship between range size (the number of meadows a species was observed at) and abundance (Appendix 1: Fig. S4). This highlights a potential bias in studies like ours against detecting dispersal limitation if it is most severe for rare taxa. Increasing the sample size might reduce the risk of this type of bias, but would cost a substantial amount of time associated with invertebrate identification. More efficient biodiversity sampling methods (e.g. eDNA) could overcome this time cost.

Theory predicts that intermediate dispersal (‘dispersal sufficiency’, Leibold and Chase 2017) allows species to colonize sites where local environmental conditions are optimal for growth and reproduction, whereas high dispersal (‘dispersal surplus’) overrides this, allowing species to persist in habitats that cannot sustain positive population growth without substantial immigration (‘mass effects’, Mouquet and Loreau 2013). A subset of species was present at all meadows, but their abundances varied (Fig 3d); this pattern may suggest weak mass effects, where dispersal rates are high enough for several taxa to occur at most meadows even if local conditions are suboptimal, but not so high as to completely overwhelm the signature of environmental conditions (Schmida and Wilson 1985, Mouquet and Loreau 2003). Metacommunity theory predicts that such weak mass effects are likely whenever dispersal rates are not limiting (Thompson et al. 2017), particularly in organisms that cannot control their own dispersal (Leibold and Chase 2017) such as seagrass-associated invertebrates. This may explain the poor model fit in some ubiquitous taxa such as *Amphissa columbiana* (R^2^ = 0.05, Appendix 1: Table S2). The poor model fit may suggest that these taxa either 1) persist at several meadows even if local environmental conditions are not optimal, due to sufficient immigration, or 2) have broader environmental niches, and thus the environmental gradient observed across the study region is not sufficient to observe clear responses (lower or higher abundances) in these taxa.

### Regional species abundance patterns suggest weak environmental filtering

We found that differences in local-scale environmental factors account for some differences in invertebrate presence and abundance (H2), however there remains a great degree of unexplained variation given the imperfect model fit (mean R^2^ = 0.27). Abiotic environmental variables had a greater influence on invertebrate distribution and abundance than biotic variables (Fig. 2a). The estimated *β* parameters suggest that, generally speaking, environments that are saltier, have a higher seagrass surface area, smaller annual temperature range, and higher dissolved oxygen levels have higher abundances of many taxa in this study (Fig. 2b). However, there were several exceptions to this trend, highlighting the importance of understanding individual species responses to the environment as opposed to the community as a whole. Overall, we conclude that, at least in taxa for whom the model that had a higher fit (e.g. *Leptochelia* sp., R^2^ = 0.52), environmental filtering influences regional patterns in abundance and distribution.

Differences in environmental conditions may influence distribution and abundance across the region in one of three ways. First, abiotic variables such as temperature and nutrient availability may influence food availability through primary productivity. This is possible despite the fact that biometric variables (seagrass LAI, biomass) explained a relatively low proportion of variance in abundance; while these measurements were taken at a single time point during field sampling, the abiotic variables represented annual averages (Assis et al. 2018) and therefore better represent long-term productivity. Second, temperature and salinity may affect species’ abundances via environmental tolerance ranges; for example, the isopod *Idotea baltica* (a relative of *P. resecata* and *P. montereyensis* in our study) experiences significantly lower survival in prolonged exposure to higher temperatures and lower salinities than those in their home habitat (Rugiu et al. 2018). Third, temperature may influence invertebrate metabolic demands, thereby influencing consumption rates and available primary biomass (O’Connor 2009). Reduced food availability may increase competition, thus altering the number of individuals a seagrass patch can host.

Environmental conditions in temperate seagrass meadows fluctuate seasonally, driving changes in epifaunal community structure throughout the year (Nelson et al. 1979, Wlodarska-Kowalczuk et al. 2014, Whippo et al. 2018). Sampling meadows several times within a year to capture these temporal changes would clarify the importance of the environment if invertebrate assemblages shift with changes in environmental variables following the linear relationships uncovered in our analysis.

### Species co-occurrence patterns may suggest the importance of interspecific interactions

We did not observe strong between-region co-occurrence patterns, suggesting that there do not appear to be clear regional assemblages corresponding with the five subregions in our analysis (Figs. 3c, black symbols in 3d). We also did not observe strong co-occurrence patterns between quadrats, however we did observe strong co-occurrence patterns at the meadow-level spatial scale. The explanation is that meadows had distinct assemblages, and these assemblages tended to be homogenized within meadows; if two taxa co-occurred in a given meadow (Fig. 3b), then they likely co-occurred in all quadrats within the meadow (Fig. 3a). This is consistent with previous findings that epifaunal diversity patterns do not differ from random patterns within meadows (between quadrats) (Whippo et al. 2018).

At the meadow-level, two main co-occurrence groupings showed antagonistic abundance patterns across meadows that could not be explained by spatial structure or shared responses to the environment (Fig. 3d). These co-occurrence groups have not, to our knowledge, been explicitly documented before in this region. While there are multiple possible explanations for non-random species co-occurrence patterns (Connor and Simberloff 1979), we ruled out distance between meadows and measured environmental variables, because co-occurrence values were extracted from residual variation unexplained by the environment or space (Ovaskainen et al. 2017). This residual variation therefore likely can be explained by a combination of unmeasured environmental variables, stochasticity, and biotic interactions.

Several interaction types may influence abundance and diversity. The majority of species in our survey are herbivores or detritivores, and thus may compete for primary (mostly epiphytic algae) biomass. Laboratory experiments have shown that grazing rate and habitat selection in amphipods are altered in the presence of morphologically and functionally similar interspecific competitors (Howard 1985, Brooks and Bell 2001, Beermann et al. 2018). Other species in our study are suspension feeders, and thus may not compete for food but possibly predator-free space or substrate on eelgrass blades.

The invertebrate assemblage at a given meadow may have multiple possible compositional states depending on the arrival order of species; this phenomenon is known as priority effects (Fukami et al. 2016, Ke and Letten 2018) and has been documented in marine fouling communities (Vieira et al. 2018). The antagonistic co-occurrence groupings in our analysis could suggest priority effects at the meadow-level. It is possible that, following seasonal declines in abundance or a disturbance event, the first few populations to increase in abundance or colonize a meadow determine the success of others. Abundance patterns in the sessile calcifying polychaete *Spirorbis* sp. might demonstrate an example of priority effects. *Spirorbis* sp. dominated the HL meadow on Haida Gwaii; there were approximately 16 300 individuals, several orders of magnitude higher than its abundance elsewhere. Meadow HL also had the lowest taxonomic richness of all 17 meadows (9 species), and we observed fewer micro and macro-epiphytes on eelgrass blades. Instead, the eelgrass was completely covered with *Spirorbis* sp. This phenomenon has also been observed in *Z. marina* meadows in Akkeshi-Ko estuary, Japan (Clark et al. 2018) and in *Thalassia testudinum* meadows in the Northwestern Gulf of Mexico (Dirnberger 1990). Experimental evidence suggests that *Spirorbis* spp. larvae tend to settle lower in the water column on newer seagrass growth, away from epiphytic algae and previously settled conspecifics (Dirnberger 1990). Settlement rates were determined by planktonic larval density rather than space availability on seagrass blades (Dirnberger 1990). Given this, it is possible that *Spirorbis* sp. dominates HL from a combination of density-dependent processes (high larval recruitment) and environmental conditions (high salinity, low nitrates). The low epiphyte load on seagrass at HL, whether mostly driven by high *Spirorbis* sp. densities or by the environment, may explain the low abundance and diversity of all other invertebrate taxa. Overall, the importance of priority effects likely depends on the age, colonization history, and disturbance regime of seagrass meadows.

Negative co-occurrence patterns in taxonomic distributions could also reflect the higher abundance of predators at some meadows. Field experiments have shown that changes in predation pressure by fish, shorebirds, and predatory invertebrates can shift seagrass-associated epifaunal assemblages in a matter of weeks (Amundrud et al. 2015; Huang et al. 2015). Previous studies involving some of the meadows in this analysis (Haida Gwaii, Clayoquot Sound, and Barkley Sound) found high variation (beta diversity) in fish assemblages among meadows (Robinson et al. 2011, Iacarella et al. 2018). A comprehensive survey of fish and bird predators is required to determine the extent to which top-down trophic interactions structure these invertebrate communities. Positive co-occurrence patterns may additionally be a result of positive biotic interactions. An example is *Orchomenella recondita* - a gammarid amphipod that lives in the gastrovascular cavity of the anemone *Anthopleura elegans* (Carlton 2007). This species was only recorded at the SS meadow, and specifically only found in quadrats where *A. elegans* was collected with the seagrass shoots.

While inferences from data taken at a single time point have limitations, our JDSM approach suggests that there is more to learn about species interactions and the interplay between interactions and dispersal in structuring these communities. The hypothesized processes driving seagrass epifaunal diversity patterns that can be further tested with experiments to test for priority effects or trophic interactions, with population genetics to test for demographic connectivity, or with particle tracking models (e.g. Treml et al. 2008) to estimate dispersal velocity.

## Conclusion

Seagrasses are important coastal foundation species, valued throughout the world for supporting diverse and productive food webs (Constanza et al. 1997, Williams and Heck 2001). The extent of these ecosystems is declining at rates believed to exceed those of coral reef and rainforest habitat loss, suggesting major losses of associated biodiversity (Waycott et al. 2009). Our finding that several seagrass-associated faunal taxa across functional groups are consistently present across a fairly large spatial scale is consistent with notion that “the mobile, fast-reproducing, and generally omnipresent animal community is keenly responsive to the presence of habitat” (Lefcheck et al. 2016). Our study suggests that the management of seagrass for its ecosystem services may be more effective if both local and regional processes are considered explicitly in habitat protection plans (Economo 2011).

## Data accessibility

Data will be made permanently available on Dryad should the manuscript be accepted. All data and code for the analysis can be currently found at https://github.com/keilast/HMS-Seagrass.

## Competing interests

We have no competing interests.

## Acknowledgements

KAS and MIO conceived of the study. JY provided samples from IN, DK, and EB. LL provided samples from RA and HL. EMA and MHL provided samples from TN, TB, CI, and CS. KAS collected and processed samples, analyzed data, and wrote the first manuscript draft. PLT provided recommendations for the analysis. KAS, PLT, and MIO contributed to writing the manuscript. This research was financially supported by a UBC Zoology SURE Grant, and NSERC Discovery Grants, Canadian Foundation for Innovation (CFI), and is sponsored by the NSERC Canadian Healthy Oceans Network and its Partners: Department of Fisheries and Oceans Canada and INREST (representing the Port of Sept-Îles and City of Sept-Îles). PLT is supported by NSERC and Killam postdoctoral fellowships. We are grateful to the Tula Foundation for providing logistical support and facilities during Calvert Island field collections. Special thanks go to Michelle Paleczny and Gulf Islands National Park Reserve for boat and laboratory space, and staff, students, and volunteers from Pacific Rim National Park Reserve for their assistance collecting eelgrass. We thank Minako Ito, Ariane Comeau, Rob Underhill, Abbie Sherwood, Giordano Bua for assistance with samples and Coreen Forbes, Matt Whalen, and anonymous reviewers for feedback on the manuscript.

## Notes

### Competing Interest Statement

The authors have declared no competing interest.

### Summary of Updates

Revised figures and minor edits to text.

## Literature Cited

Aivelo, T. and Norberg, A. 2018. Parasite-microbiota interactions potentially affect intestinal communities in wild mammals. J. Anim. Ecol. 18:438–447.

Amundrud, S. L. et al. 2015. Indirect effects of predators control herbivore richness and abundance in a benthic eelgrass (*Zostera marina*) mesograzer community. – J. Anim. Biol. 84: 1092–1102.

Assis, J. et al. 2018. Bio-ORACLE v2.0: Extending marine data layers for bioclimatic modelling. – Global Ecol. Biogeogr. 27:277–284.

Beermann, J. et al. 2018. Combined effects of predator cues and competition define habitat choice and food consumption of amphipod mesograzers. - Oecologia 186:645–654.

Best, R., and Stachowicz, J. J. 2014. Phenotypic and phylogenetic evidence for the role of food and habitat in the assembly of communities of marine amphipods. - Ecolog. 95: 775–786.

Berlow, E. L. 1999. Strong effects of weak interactions in ecological communities. - Nature 398: 330–334.

Boström, C. et al. 2006. Seagrass landscapes and their effects on associated fauna: A review. - Estuarine, Coastal Shelf Sci. 68: 383–403.

Boström, C. et al. 2010. Invertebrate dispersal and habitat heterogeneity: Expression of biological traits in a seagrass landscape. J. Exp. Mar. Biol. Ecol. 390:106–117.

Bray, J. R. and J. T. Curtis. 1957. An ordination of upland forest communities of southern Wisconsin. – Ecol. Monogr. 27:325–349.

Brooks R.A., and Bell S.S. 2001. Mobile corridors in marine landscapes: Enhancement of faunal exchange at seagrass/sand ecotones. 2001. – J. Exp. Mar. Bio. Ecol. 264: 67–84.

Brown, B. L. et al. 2017. Making sense of metacommunities: dispelling the mythology of a metacommunity typology. - Oecologia 183:643–652.

Carlton, J. 2007. The Light and Smith Manual: Intertidal invertebrates from Southern California to Oregon. Fourth edition. University of California Press, Oakland, California, USA.

Cébrian, J. and Duarte, C. M. 1998. Patterns in leaf herbivory on seagrasses. – Aquat. Bot. 60:67–82.

Connor, E. F. and Simberloff, D. 1979. The assembly of species communities: Chance or competition? - Ecology 60:1132–1140.

Costanza, R., et al. 1997. The value of the world’s ecosystem services and natural capital. - Nature 387:253–260.

Cottenie, K. 2005. Integrating environmental and spatial processes in ecological community dynamics. – Ecol. Lett. 8:1175–1182.

Duffy J. E. et al. 2015. Biodiversity mediates top-down control in eelgrass ecosystems: A global comparative-experimental approach. – Ecol. Lett. 18:696–705.

Economo, E. P. 2011. Biodiversity conservation in metacommunity networks: Linking pattern and persistence. – Amer. Nat. 177:167–180.

Fukami, T. et al. 2016. A framework for priority effects. – J. Veg. Sci. 27:655–657.

Gaines, S. and Roughgarden, J. 1985. Larval settlement rate: A leading determinant of structure in an ecological community of the marine intertidal zone. – Proc. Natl. Acad. Sci. 82:3707–-3711.

Gilbert, B. and Bennett, J. R. 2010. Partitioning variation in ecological communities: do the numbers add up? – J. Appl. Ecol. 47:1071–1082.

Guillaume Blanchet, F. et al. 2018. HMSC: Hierarchical Modelling of Species Community. R package version. 2.1-2.

Gullström, M. et al. 2012. Spatial patterns and environmental correlates in leaf-associated epifaunal assemblages of temperate seagrass (*Zostera marina*) meadows. – Mar. Biol. 159:413–425.

Huang, A. C. et al. 2015. Top-down control by great blue herons *Ardea herodias* regulates seagrass-associated epifauna. - Oikos 124:1492–1501.

Howard, R. K. 1985. Measurements of short-term turnover of epifauna within seagrass beds using an in situ staining method. – Mar. Ecol. Prog. Ser. 22: 163–168.

Ke, P. J. and Letten, A. D. 2018. Coexistence theory and the frequency-dependence of priority effects. – Nat. Ecol. Evol. 2:1691–1695.1–7.

Kinlan, B. P. and Gaines, S. D. 2003. Propagule dispersal in marine and terrestrial environments: A community perspective. - Ecology 84:2007–2020.

Kozloff, E. 1996. Marine Invertebrates of the Pacific Northwest. University of Washington Press, Seattle, Washington, USA.

Leibold, M. A. et al. 2004. The metacommunity concept: A framework for multi-scale community ecology. – Ecol. Lett. 7:601–613.

Leibold, M. A. and Chase, J. M. 2017. Metacommunity Ecology, Volume 59. Princeton University Press, Princeton, New Jersey, USA.

Lefcheck, J. S. et al. 2016. Faunal Communities Are Invariant to Fragmentation in Experimental Seagrass Landscapes. - PLoS ONE 11(5): e0156550.

Levin, S. A. and Paine, R. T. 1974. Disturbance, patch formation, and community structure. – Proc. Natl. Acad. Sci. 71:2744–2747.

Logue, J. et al. 2011. Empirical approaches to metacommunities: a review and comparison with theory. – Trends Ecol. Evol. 26:482–491.

Mitarai, S. D. et al. 2008. A numerical study of stochastic larval settlement in the California Current system. – J. Marine Syst. 69:295–309.

Morlon, H. et al. 2008. A general framework for the distance-decay of similarity in ecological communities. - Ecol. Lett. 11:904–917.

Mouquet, N. and Loreau, M. 2003. Community patterns in source-sink metacommunities. – Amer. Nat. 162:554–557.

Nelson, W. G. 1979. An analysis of structural pattern in an eelgrass (*Zostera marina* L.) amphipod community. - J. exp. Mar. Biol. Ecol. 39:231–264.

O’Connor, M. I. 2009. Warming strengthens an herbivore-plant interaction. - Ecology 90:388–398.

Orth, R. J. et al. 1984. Faunal communities in seagrass beds: A review of the influence of plant structure and prey characteristics on predator-prey relationships. - Estuaries 7:339–350.

Ovaskainen, O. et al. 2016a. Uncovering hidden spatial structure in species communities with spatially explicit joint species distribution models. - Methods Ecol. Evol. 7:428–436.

Ovaskainen, O. et al. 2016b. Using latent variable models to identify large networks of species-to-species associations at different spatial scales. - Methods Ecol. Evol. 7:549–555.

Ovaskainen, O. et al. 2017. How to make more out of community data? A conceptual framework and its implementation as models and software. – Ecol. Lett. 20:561–576.

R Development Core Team. 2014. R: a language and environment for statistical computing. R Foundation for Statistical Computing. Vienna, Austria. http://www.R-project.org.

Ricklefs, R. E. and Schluter, D. 1993. Species Diversity in Ecological Communities: Historical and Geographical Perspectives. University of Chicago Press, Chicago, Illinois, USA.

Rugiu, L. et al. 2018. Variations in tolerance to climate change in a key littoral herbivore. – Mar. Biol. 165:1–11.

Sala, E. and Graham, M. H. 2002. Community-wide distribution of predator-prey interaction strength in kelp forests. – Proc. Natl. Acad. Sci. 99:3678–3683.

Sand-Jensen. 1977. Effect of epiphytes on eelgrass photosynthesis. – Aquat. Bot. 3:55–63.

Schmida, A. and Wilson, M. V. 1985. Biological determinants of species diversity. – J. Biogeogr. 12:1–20.

Stier, A. C. et al. 2019. Temporal variation in dispersal modifies dispersal-diversity relationships in an experimental seagrass metacommunity. – Mar. Ecol. Prog. Ser. 613:67–76.

Thom, R. et al. 1995. Temporal patterns of grazers and vegetation in a temperate seagrass system. – Aquat. Bot. 50:201–205.

Thompson, P. L. et al. 2017. Loss of habitat and connectivity erodes species diversity, ecosystem functioning, and stability in metacommunity networks. - Ecography 40:98–108.

Thompson, P. L. et al. 2020. A process-based metacommunity framework linking local and regional scale community ecology. – Ecol. Lett. (In press)

Treml, E. A. et al. 2008. Modeling population connectivity by ocean currents: a graph-theoretic approach for marine conservation. – Landsc. Ecol. 23:19–36.

Tuomisto, H. et al. 2012. Modelling niche and neutral dynamics: on the ecological interpretation of variation partitioning. - Ecography 35:961–971.

Tyberghein, L. et al. 2011. Bio-ORACLE: a global environmental dataset for marine species distribution modelling. – Glob. Ecol. Biogeogr. 21: 272–281.

Vieira, E. et al. 2018. Persistence and space preemption explain species-specific founder effects on the organization of marine sessile communities. - Ecology and Evolution 8:3430–3442.

Virnstein, R. W., and R. K. Howard. 1987. Motile epifauna of marine macrophytes in the Indian River Lagoon, Florida. II. Comparisons between drift algae and three species of seagrasses. Bulletin of Marine Science 41:13–26.

Vellend, M. 2010. Conceptual synthesis in community ecology. – Q. Rev. Biol. 85:183–206.

Ward, J. H. 1963. Hierarchical grouping to optimize an objective ounction. - Journal of the American Statistical Association 58:236–244.

Waycott, M. et al. 2009. Accelerating loss of seagrasses across the globe threatens coastal ecosystems. – Proc. Natl. Acad. Sci. 106: 12377–12381.

Wichmann, C. S. et al. 2012. Floating kelps in Patagonian Fjords: An important vehicle for rafting invertebrates and its relevance for biogeography. – Mar. Biol. 159:2035–2049.

Williams, S. L., and K. L. Heck, Jr. 2001. Seagrass community ecology. Pages 317–337 in M. D. Bertness, M. E. Hay, and S. D. Gaines, editors. Marine community ecology. Sinauer Associates, Sunderland, Massachusetts, USA.

Whalen, M. A., et al. 2013. Temporal shifts in top-down vs. bottom-up control of epiphytic algae in a seagrass ecosystem. - Ecology 94:510–520.

Whippo, R. et al. 2018. Epifaunal diversity patterns within and among seagrass meadows suggest landscape-scale biodiversity processes. - Ecosphere 9:02490.10.1002/ecs2.2490.

Wlodarska-Kowalczuk, M. et al. 2014. Evidence of season-dependency in vegetation effects on macrofauna in temperate seagrass meadows (Baltic Sea). - PLoS One 9:e100788.

Yamada et al. 2007. Temporal and spatial macrofaunal community changes along a salinity gradient in seagrass meadows of Akkeshi-ko estuary and Akkeshi Bay, northern Japan. – Hydrobiologia 592: 345–358.

Yamada, K. et al. 2014. Environmental and spatial controls of macroinvertebrate functional assemblages in seagrass ecosystems along the Pacific coast of northern Japan. - Glob. Ecol. Cons. 2:47–61.

Yeager, L. A. et al. 2019. Trait sensitivities to seagrass fragmentation across spatial scales shape benthic community structure. – J. Anim. Ecol. 88:1743–1754.

